## supplemental fig 1 and 2 for "Multiple Sclerosis Patients have an Altered Gut Mycobiome and Increased Fungal to Bacterial Richness"

### Supporting Information

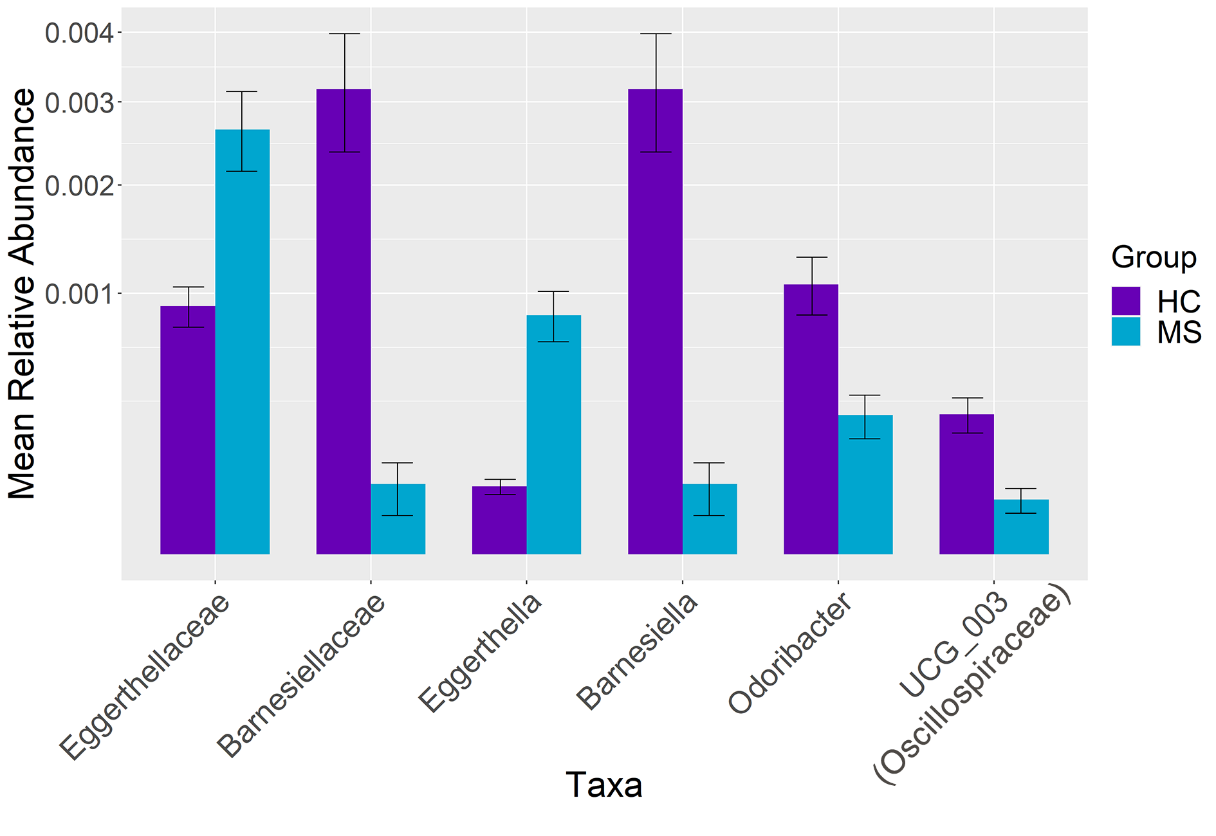

### S1 Fig. Relative Abundance of Differentially Abundant Families and Genera in HC and MS Bar plot showing relative abundances of differentially abundant taxa (p < 0.05) at the family and genus level

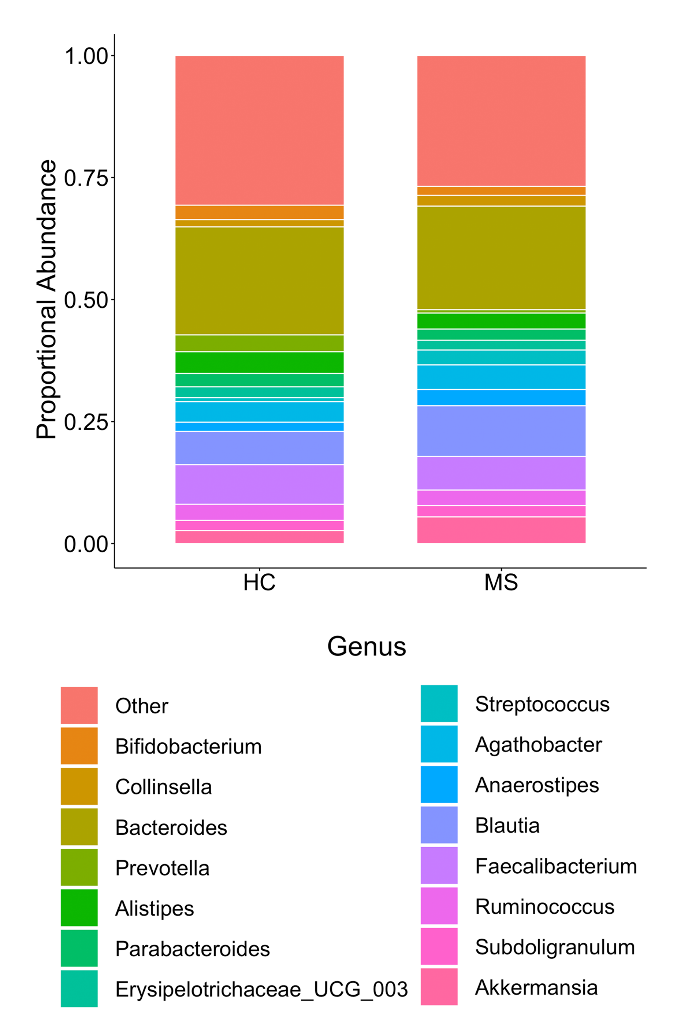

### S2 Fig. Proportional Abundance of Bacteria at the Genus Level Stacked bar plots representing the proprotional abundance of the top 10 bacteria at the genus level in MS and HC groups
